# Microvascular pathology of proteotoxic endothelial signature characterizes Progressive Supranuclear Palsy

**DOI:** 10.64898/2026.07.16.738175

**Authors:** Seojin Lee, Xia Han, Satoshi Tanakawa, Takeru Kuwabara, Koji Yoshida, Shelley L. Forrest, Shojiro Ichimata, Hidetomo Tanaka, Tomoya Kon, Shinya Tanaka, Ekatarina Rogaeva, Maria Carmela Tartaglia, Susan H. Fox, Anthony E. Lang, Jessica E. Rexach, Gabor G. Kovacs

## Abstract

Cerebrovascular pathology is increasingly implicated in neurodegenerative diseases, yet its pathomechanistic contribution remains poorly defined. Building on prior evidence of dysregulated iron and oxygen homeostasis in early-affected brain regions of progressive supranuclear palsy (PSP), we hypothesized that brain microvascular alterations may play an etiological role in select neurodegenerative proteinopathies. First, we conducted a systematic neuropathological evaluation of 178 brains from the University Health Network Neurodegenerative Brain Collection, including Alzheimer’s disease-related neuropathologic change (ADNC; n=30), Lewy body disease with high or intermediate ADNC (n=38) and low ADNC (n=16), multiple system atrophy (MSA; n=14), PSP (n=39), frontotemporal lobar degeneration with TDP-43 proteinopathy (FTLD-TDP; n=10), and controls (n=31). Arteriolosclerosis, microinfarction, and calcification were assessed in the basal ganglia and frontal cortex. Iron burden was correlated by quantification of Perl’s staining in MSA and PSP, where vessel pathology was most severe. Single-nucleus RNA-sequencing (snRNA-seq) of frontal cortex tissue from control (n=5) and PSP (n=8) cases with varying arteriolosclerosis severity was performed to characterize the vascular transcriptome, with validation against an independent snRNA-seq evaluation of PSP (n= 11), Pick’s disease (n=9), AD (n=10), and control (n=10) brains. Histological analysis revealed disease-specific involvement of microvascular pathology in neurodegenerative diseases, identifying PSP to demonstrate most prominent and widespread vessel wall thickening across regions examined. Regression analysis using demographic, *APOE* and *MAPT* genetic risk status, and neuropathological features of cases corroborated the distinct association with PSP pathology. Elevated iron load in early affected regions of MSA and PSP brains correlated with greater vessel wall thickening, suggesting a possible pathomechanistic relationship between the two disease physiologies. snRNA-seq analysis of vascular transcriptome identified robust upregulation of heat shock proteins and hypoxia-related genes in PSP endothelial cells and pericytes across both datasets. Importantly, we found the proteotoxic signature to be strongly associated with higher vessel scores in PSP cases, linking microvascular morphology to endothelial dysfunction. Our comprehensive neuropathological evaluation coupled with correlative snRNA-seq analysis establish PSP-specific arteriolar thickening associated with endothelial proteotoxic state as a candidate pathogenic mechanism. The cerebral arteriolar unit represents a compelling therapeutic target for disease modification in PSP.

## Introduction

Neurodegenerative diseases are characterized by the accumulation of misfolded proteins in the brain, which is associated with synaptic dysfunction and widespread neuronal and glial network impairment.^1^ Beyond the frequent coexistence of multiple neurodegenerative proteinopathies, cerebrovascular co-pathology is commonly observed^2^ and thus the involvement of vascular changes in the pathophysiology of neurodegenerative diseases are increasingly considered. Nevertheless, vascular lesions are markedly heterogeneous in type, location, and severity, and their underlying mechanisms and relative contributions—alone or in combination—to neurodegenerative disease pathogenesis remain unclear.

Difficulties delineating the vascular-associated disease pathomechanisms are in part due to limitations in both clinical and neuropathological approaches used to assess vascular co–pathologies in patient cohorts and tissue samples. White matter hyperintensity (WMH), a widely used MRI marker of cerebral small vessel disease (SVD), is increased in several neurodegenerative disorders, including sporadic behavioural variant frontotemporal dementia (bvFTD), multiple system atrophy (MSA), Parkinson’s disease (PD), and progressive supranuclear palsy (PSP).^3–7^ Further, cerebrovascular alterations such as enlarged perivascular spaces, lacunes, arterial wall thickening, and carotid stenosis evaluated using MRI in patients of PSP and MSA have been shown to correlate with disease progression and clinical outcomes.^7,8^ However, clinical neuroimaging primarily captures global or regional alterations at a resolution that is susceptible to misinterpretation and unlikely to detect aging-related SVD or subtle vascular lesions. Consequently, a comprehensive understanding of vascular contributions to neurodegenerative disease requires detailed microscopic assessment of vascular changes in post-mortem tissue. Neuropathological findings, however, are strongly influenced by heterogenous assessment methodologies which vary by evaluation of single or multiple lesion types and regions, as well as subjective interpretation. The introduction of the Vascular Cognitive Impairment Neuropathology Guidelines (VCING) has improved inter-center consistency by emphasizing specific lesion types within defined brain regions, particularly occipital vessel pathology and basal ganglia microinfarcts.^9^ Yet holistic interpretation of existing neuropathological reports remains challenging, and systematic regional mapping of SVD remains limited – especially beyond Alzheimer’s disease (AD) and Lewy body disease (LBD), for which findings remain inconsistent.^10–14^

Currently, cerebrovascular pathology is commonly thought to follow protein pathology. However, select neuropathological studies have reported observations that are inconsistent with this model. AD and LBD brains with concomitant cerebrovascular lesions have shown lower burdens of their respective lesions than those without cerebrovascular disease.^15^ Similarly, negative correlations were reported between cerebrovascular lesions and tau, TDP-43, and Aβ.^16^ While these observations have been interpreted as evidence that concomitant cerebrovascular lesions lower the threshold for dementia, they also raise the possibility that proteinopathy–related vascular alterations do not always occur secondary to pathological protein burden. Further supporting the possibility of early involvement of vascular dysfunction in disease, neuropathological alterations associated with hypoxia – which results from an impairment in the cerebral arteriolar vascular unit (cAVU) comprised of endothelial cells (EC), pericytes (PC), smooth muscle cells, inflammatory cells, fibroblast-like cells, and astrocytic end-feet^17^ – is commonly observed with aging and in early stages of neurodegenerative diseases.^18,19^ Moreover, brain arteriolosclerosis has been correlated with aging related tau astrogliopathy (ARTAG)^20^, and importantly, accumulation of astrocytic tau pathology in ARTAG and PSP show early selective vulnerability to perivascular areas.^21,22^

Therefore, we hypothesize that SVD contributes early to the pathogenesis of select neurodegenerative diseases. A comparative study across multiple neurodegenerative diseases reported pronounced SVD involvement in Pick’s disease (PiD), with no major differences among AD, PSP, corticobasal degeneration (CBD), and argyrophilic grain disease (AGD); however, this analysis was based on the extent of regional involvement across the brain rather than quantitative severity measures.^23^ Therefore, comparisons of SVD severity in neurodegenerative diseases are lacking. We performed a systematic neuropathological assessment of arteriolar wall thickening, microinfarction, and vascular calcification in post-mortem brains of neurodegenerative proteinopathies, including AD, AD with LBD (AD+LBD), LBD, MSA, PSP, and frontotemporal lobar degeneration with TDP–43 proteinopathy (FTLD–TDP). Finding a marked microvascular phenotype in PSP, we interrogated underlying mechanisms using single-nucleus RNA sequencing (snRNA-seq) analyses of vascular cells from two independent datasets.

## Materials and Methods

### Case selection

A retrospective analysis of 178 neuropathological autopsies from the University Health Network Neurodegenerative Brain Collection (UHN-NBC) was conducted. All brain samples were obtained during autopsies following appropriate consenting procedures and with approval from the Local Ethical Committee, the UHN Research Ethics Board (Nr. 20-5258), and the University of Toronto (Nr. 39459). The autopsies were performed in accordance with the ethical standards outlined in the 1964 Declaration of Helsinki, updated in 2008.

Following diagnostic staining of brain sections (Supplementary methods), the assessment of AD pathology was based on the Braak NFT stage,^24^ Thal phase,^25^ and CERAD (Consortium to Establish a Registry for Alzheimer’s Disease) criteria.^26^ The level of Alzheimer’s disease neuropathologic change (ADNC) was determined using the ABC score as per the guidelines set by the National Institute on Aging -Alzheimer’s Association (NIAAA).^27^ The severity of Lewy-related pathology was evaluated following the consensus criteria for LBD using a-syn immunohistochemistry.^28^ Cases with intermediate and high ADNC levels were classified as AD, those with low ADNC and limbic or neocortical Lewy-related pathology as LBD, and those meeting both criteria as AD+LBD. PSP was diagnosed based on established criteria^29^ and staging was performed as previously reported^30^; AGD was classified according to the Saito stage^31^; and TDP-43 pathology was categorized either as FTLD-TDP^32^ or as limbic-predominant age-related TDP-43 encephalopathy neuropathological changes (LATE-NC) stage.^33^

The cohort comprised 30 AD, 38 AD+LBD, 16 LBD, 39 PSP, 14 MSA, 10 FTLD-TDP, and 31 controls (Table 1). Controls were defined as cases with either an absence or low ADNC and no other major degenerative diseases. In 93 out of the total 178 cases for which DNA samples were available, genotypes of apolipoprotein E (*APOE*) and microtubule-associated protein tau (*MAPT*) were examined as described previously.^34^ Genotypes for each disease group are summarized in Table 1.

**Table 1.** Case Summary.

|  | Control (n=31) | AD (n=30) | AD+LBD (n=38) | LBD (n=16) | MSA (n=14) | PSP (n=39) | FTLD-TDP (n=10) |
| --- | --- | --- | --- | --- | --- | --- | --- |
| **Age, mean ± SD, years** | 64.7 ± 15.8 | 76.0 ± 11.0 | 75.2 ± 9.4 | 76.5 ± 8.3 | 66.8 ± 8.4 | 76.2 ± 7.3 | 68.1 ± 13.5 |
| **Sex (female/male)** | 11/20 | 19/11 | 20/18 | 5/11 | 7/7 | 17/22 | 5/5 |
| **ADNC level, mean ± SD** | 0.5 ± 0.5 | 2.9 ± 0.3 | 2.5 ± 0.5 | 0.8 ± 0.4 | 0.4 ± 0.5 | 0.9 ± 0.8 | 0.5 ± 0.7 |
| **A Score, mean ± SD** | 0.7 ± 0.8 | 3.0 ± 0.0 | 2.6 ± 0.5 | 1.2 ± 0.9 | 0.6 ± 0.9 | 1.1 ± 1.1 | 0.4 ± 0.5 |
| **B Score, mean ± SD** | 0.6 ± 0.5 | 2.9 ± 0.3 | 2.7 ± 0.5 | 1.2 ± 0.4 | 1.0 ± 0.4 | 1.4 ± 0.6 | 1.2 ± 0.4 |
| **C Score, mean ± SD** | 0.6 ± 0.8 | 2.9 ± 0.3 | 2.6 ± 0.6 | 0.9 ± 0.8 | 0.3 ± 0.6 | 0.9 ± 1.0 | 0.7 ± 1.1 |
| **Lewy related pathology positive, n (%)** | 0 (0.0) | 13 (43.3) | 38 (100.0) | 16 (100.0) | 0 (0.0) | 11 (28.2) | 1 (10.0) |
| **LATE-NC positive, n (%)** | 0 (0.0) | 8 (26.7) | 20 (52.6) | 3 (18.8) | 1 (7.1) | 6 (15.4) | 10 (100.0) |
| **AGD positive, n (%)** | 2 (6.5) | 5 (16.1) | 6 (15.8) | 8 (50.0) | 3 (21.4) | 17 (44.6) | 1 (10.0) |
| **PSP stage, n ≥4, (%)** | 0 (0.0) | 0 (0.0) | 0 (0.0) | 0 (0.0) | 0 (0.0) | 30 (81.1) | 0 (0.0) |
| **APOE gene** |  |  |  |  |  |  |  |
| **ε2/ε2, n (%)** | 0 (0.0) | 0 (0.0) | 0 (0.0) | 0 (0.0) | 0 (0.0) | 0 (0.0) | 1 (25.0) |
| **ε2/ε3, n (%)** | 0 (0.0) | 0 (0.0) | 1 (3.2) | 0 (0.0) | 0 (0.0) | 2 (8.0) | 0 (0.0) |
| **ε2/ε4, n (%)** | 0 (0.0) | 0 (0.0) | 0 (0.0) | 1 (8.3) | 1 (7.7) | 3 (12.0) | 0 (0.0) |
| **ε3/ε3, n (%)** | 1 (50.0) | 5 (85.7) | 6 (19.4) | 8 (66.7) | 9 (69.2) | 13 (52.0) | 3 (75.0) |
| **ε3/ε4, n (%)** | 1 (50.0) | 1 (14.3) | 20 (64.5) | 3 (25.0) | 3 (23.1) | 6 (24.0) | 0 (0.0) |
| **ε4/ε4, n (%)** | 0 (0.0) | 0 (0.0) | 4 (12.9) | 0 (0.0) | 0 (0.0) | 1 (4.0) | 0 (0.0) |
| **MAPT gene haplotype** |  |  |  |  |  |  |  |
| **H1/H1, n (%)** | 2 (100.0) | 5 (71.4) | 15 (48.4) | 9 (75.0) | 11 (84.6) | 23 (92.0) | 3 (75.0) |
| **H1/H2, n (%)** | 0 (0.0) | 1 (14.3) | 14 (45.2) | 2 (16.7) | 1 (7.7) | 2 (8.0) | 1 (25.0) |
| **H2/H2, n (%)** | 0 (0.0) | 1 (14.3) | 2 (6.5) | 1 (8.3) | 1 (7.7) | 0 (0.0) | 0 (0.0) |

**Abbreviation:** Alzheimer's disease (AD); Lewy body disease (LBD); Multiple system atrophy (MSA); Progressive supranuclear palsy (PSP); Frontotemporal lobar degeneration with TDP-43-immunoreactive inclusions (FTLD-TDP); Standard deviation (SD); Alzheimer's disease neuropathological change (ADNC); Limbic-predominant age-related TDP-43 encephalopathy neuropathologic change (LATE-NC); Argypophilic Grain Disease (AGD); Apolipoprotein E (APOE); Microtubule-associated protein tau (MAPT).

### Evaluation of small vessel pathology

To evaluate vessel wall thickening we applied a Vessel Score, which is a modification of the VCING.^9^ Our assessment focused on vessels in the basal ganglia (BG) and the frontal lobe (FC) as these regions show greater involvement of protein pathology across our disease cohort than the occipital lobe which was the focus of the VCING evaluation strategy. The scores were rated on a scale from 0 to 3 (Supplementary Fig. 1A-D) for each region (BG and FC separately) and then combined to generate a total score. Additionally, the presence of vascular calcification (Supplementary Fig. 1E) and microinfarcts (Supplementary Fig. 1F) were assessed separately in the BG and FC. The evaluations were blindly confirmed by an independent neuropathologist.

### Quantification of iron load

4.5µm-thick FFPE tissue sections of PSP (n=35) and MSA (n= 12) posterior BG were stained using Perl’s iron staining and staining positivity in the globus pallidus (GP) were quantified as previously described.^35^ Briefly, sections were deparaffinized and rehydrated, then incubated in a solution of 2% potassium ferrocyanide and 2% HCI for 30 minutes followed by Nuclear Fast Red counterstain. On scanned whole-sections, GP was manually annotated and Perl’s positivity within was quantified as % annotated tissue using the HALO software (Indica Labs).

### Single-nucleus RNA sequencing analysis

#### Nuclei isolation, library preparation and data preprocessing

SnRNA-seq of frozen middle FC from PSP (n=8) and control (n=5) brains were performed as described previously.^36^ Briefly, nuclei were isolated from fresh-frozen tissue using either a Singulator 100 (S2 Genomics) or the Chromium Nuclei Isolation Kit with RNase inhibitor (10x Genomics), followed by 10x Chromium 3′ gene-expression library preparation (v3.1 or GEM-X v4) and sequencing on Illumina NovaSeq instruments (NovaSeq 6000 or NovaSeq X. Reads were processed with Cell Ranger (v9.0.1),^37^ and ambient RNA was removed using CellBender (v0.3.0).^38^ Downstream analyses were carried out in Seurat (v5.1.0)^38^: nuclei with <200 or >5,000 detected genes (nFeature_RNA) or ≥10% mitochondrial RNA (percent.mt) were excluded, data were log-normalized (LogNormalize, scale.factor = 10,000; log1p), 2,000 highly variable genes were used for PCA/UMAP^39^ and doublets were removed with DoubletFinder (v2.0.6).^40^ Per-sample objects were integrated with Harmony (IntegrateLayers, method = “HarmonyIntegration”).^41^ Cell annotation was performed by reference mapping to the Azimuth (v0.5.0)^42,43^ human cortex atlas, and Seurat clusters in which a single reference-derived cell type accounted for ≥75% of nuclei were labeled as that cell type and retained for downstream cell type–specific analyses, whereas mixed or undetermined clusters were excluded. Ambient-enriched subclusters were identified and removed following Caglayan *et al*.^44^ using DropletQC (v0.0.0.9000)^45^ nuclear fraction estimates, ambClusterFind/ambMarkFind, and enrichment of ambient markers (Fisher’s exact test with Benjamini–Hochberg–adjusted P values (FDR); clusters with FDR < 0.001 and odds ratio > 3 were discarded). Marker genes were identified for each cell type (Supplementary methods).

#### Differential expression and pathway enrichment analysis

Differential gene expression between PSP and controls was assessed using Seurat. The RNA assay was set as the default, and differential expression between PSP and controls was tested with the Wilcoxon rank-sum test (FindMarkers). P-values were adjusted for multiple testing using Bonferroni correction as implemented in Seurat (Bonferroni-adjusted P-values, p_val_adj), and genes with p_val_adj < 0.05 were considered differentially expressed and plotted (Supplementary methods). For each cell type, pathway enrichment was performed on a gene ranking derived from DESeq2 (v1.48.2).^46^ Raw UMI counts from nuclei were summed per gene and per donor (Ctrl or PSP) to generate pseudobulk profiles, and low-count genes (<10 total counts across samples) were removed. A DESeq2 model (∼ group + Age_z + Sex + Reagents) was fitted, and the signed Wald statistic for the PSP versus Ctrl contrast was used as the ranking metric (genes sorted in decreasing order). Human gene set from MSigDB^47^ C5 (GO Biological Process, GOBP)^48^ was obtained via msigdbr (v25.1.1)^49^ and used as input to fgsea (v1.34.2)^50^ (minSize = 15, maxSize = 500; default permutations). Redundant gene sets were reduced with collapsePathways, and we report the resulting main pathways. For each pathway, we report the normalized enrichment score (NES), nominal P-value, and Benjamini– Hochberg–adjusted P-value (FDR); unless stated otherwise, FDR < 0.05 was considered significant. Utilizing the FC vessel score, PSP cases were sub-grouped into high (score 2; n=5) and low (score 0-1; n=3) vessel score groups and were compared using the same methodology described.

### Independent dataset analysis of PSP endothelial cells

To validate PSP-associated gene expression changes in endothelial cells, we analyzed an independent snRNA-seq dataset comprising AD (n=10), PSP (n=11), Pick’s disease (PiD; n=9), and healthy controls (n=10) across three brain regions with distinct vulnerability (precentral gyrus, insula, and calcarine cortex).^51^ Data were processed as previously described and endothelial cells were subsequently extracted, yielding 8,085 nuclei across 40 donors (see Supplementary Methods).

For cross-dataset comparison, endothelial cells from two datasets were integrated using Seurat v5.3.1.^38^ Each dataset was normalized independently (*NormalizeData* function), and highly variable genes were identified (*FindVariableFeatures* function; nfeatures = 2000 and method = “vst”). Shared features were selected for anchor-based integration (*SelectIntegrationFeatures* function). Integration anchors were computed using canonical correlation analysis (*FindIntegrationAnchors* function), and datasets were merged to generate a corrected expression matrix (*IntegrateData* function; dims=1:30). The integrated data were scaled (*ScaleData* function), subjected to principal component analysis (*RunPCA* function; npcs=30), and visualized using uniform manifold approximation and projection (UMAP).

Differential expression analysis of the Rexach *et al*. dataset was performed using a donor-level pseudobulk strategy implemented in edgeR.^52^ Analyses were conducted both within individual brain regions (precentral gyrus, insula, calcarine cortex) and across all regions, with region included as a covariate. Lowly expressed genes were filtered out (filterByExpr function), and library size normalization (*calcNormFactors* function), and dispersion estimation (*estimateDisp* function) were performed using standard edgeR workflows. Generalized linear models were fitted to test for disease effects. In the region-specific analysis, the design formula was ∼disease + library_batch; in the region-adjusted model, the formula was ∼disease + library_batch + region. Differential expression was assessed using quasi-likelihood F-tests (*glmQLFTest* function), and significance was determined after multiple testing correction using the Benjamini-Hochberg false discovery rate (FDR). Gene Ontology and KEGG pathway enrichment analyses were performed using ShinyGO (v0.85),^53^ and terms with FDR < 0.1 were considered significant. For DEG visualization, genes were further restricted to those with robust expression, defined as log2CPM ≥ 4 in at least 4 pseudobulk samples within each disease group.

#### SCENIC regulon analysis

Gene regulatory networks were reconstructed using pySCENIC.^54^ Genes detected in at least 5% of cells were retained, and datasets were further filtered to ensure sufficient coverage (≥200 cells and ≥1,000 genes per dataset), yielding 8,237 genes from the Rexach *et al*. dataset and 7,461 genes from the Tanikawa *et al*. & Lee *et al*. dataset. Regulatory network inference was performed using the GRNBoost2 algorithm to identify transcription factor (TF)-target co-expression relationships. Co-expression modules were then derived using the *modules_from_adjacencies* function and refined by motif enrichment analysis with the *prune2df* function using hg38 cisTarget databases covering 500 bp upstream and 100 bp downstream of transcription start sites and 10 kb flanking regions, together with curated motif annotations (motifs-v10-nr). Regulons were then assembled (*df2regulons* function), and targets lacking significant motif support for the corresponding TF were pruned to generate high-confidence regulons.

Regulon activity in individual cells was quantified using AUCell, which scores the enrichment of regulon target genes within ranked gene expression profiles. To quantify condition specificity, we calculated the regulon specificity score (RSS) using the Jensen-Shannon divergence.^55^ For each regulon, AUCell activity across all cells was normalized to a probability distribution 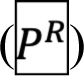, and compared to a diagnosis-specific reference distribution 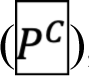, defined as a uniform distribution over cells belonging to the diagnosis of interest and zero elsewhere. Jensen-Shannon divergence was calculated as

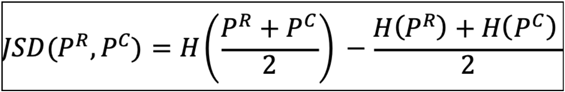

Where H denotes Shannon entropy, and RSS was defined as

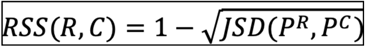

RSS values range from 0 to 1, with higher values indicating greater specificity to the given condition. RSS was computed for each regulon within diagnosis groups, excluding groups with fewer than 25 cells. For cross-dataset comparison, regulons were classified based on their relative specificity patterns as either PSP-enriched or control-enriched, and the consistency of this classification across datasets was assessed.

### Immunofluorescence of endothelial HSP90AA1 expression

FC FFPE sections of 8 PSP and2 control brains from the snRNA-seq experiment cohort (cases which have available tissue sections for immunolabelling), as well as MSA brains (n= 4) of varying FC vessel scores from 0 to 2, were labelled with claudin-5 (CLDN5; clone 4C3C2, 1:50 Invitrogen) and HSP90AA1 antibody (1:600; Invitrogen) after deparaffinization and DAKO PT-Link heat-induced antigen retrieval with low pH target retrieval solution (pH6.5; Dako). Goat anti-rabbit Alexa Fluor 488 (1:500; Invitrogen) and Donkey anti-mouse Alexa Fluor 555 (1:500; Invitrogen) was used with DAPI. Immunolabeled whole sections were scanned with Akoya PhenoImager (Akoya Biosciences). Scans were imported to HALO software and seven identical regions of interest (ROIs) (area: 0.668715µm^2^) were randomly placed within the white matter under whole-slide view and analyzed by the Object Colocalization module. Percentage (%) of total CLDN5-positive area co-positive for HSP90AA1 was computed for each ROI and averaged for each case.

### Statistical analysis

Mean vessel scores and HSP90AA1 expression (%HSP90AA1(+)CLDN5(+)area) across control and disease groups were compared by Kruskall-Wallis test followed by Dunn’s multiple comparisons test. Frequency of microinfarct and calcification were compared by Fisher’s exact test. Correlation between iron load and vessel scores in the MSA and PSP groups were evaluated by computing two-tailed Spearman’s *r* correlations. Significance was determined by *p*<0.05. Multiple logistic regression analysis was performed to evaluate the odds ratio (OR) of high vessel scores (≥2 for BG and FC, ≥4 for Total score) by age, sex, and presence of AD (low-high ADNC), LBD, MSA, PSP, AGD, and TDP-43 pathology, regardless of primary neuropathological diagnosis of cases. Effect of H1/H1 *MAPT* haplotype and *APOE ε4* genetic status on vessel pathologies were evaluated separately.

## Results

### Vascular pathology is prominent in Progressive Supranuclear Palsy

Vessel scores were generally higher, though not uniformly significant, in both BG and FC in all neurodegenerative disease groups compared to controls (Fig. 1), and the relative anatomical involvement (BG vs FC) followed the disease-specific spatiotemporal patterns in the deposition of protein pathology (Supplementary Fig.2). Comparing the severity of vessel wall thickening in the BG, MSA showed highest vessel score (mean = 2.50±0.17), closely followed by PSP (2.39±0.10) then LBD (2.31±0.15) which were significantly higher compared to that of controls (1.39±0.18, *p*<0.05) (Fig. 1A). AD, AD+LBD, and FTLD-TDP groups did not show significant differences compared to controls. In the FC, mean vessel score was highest in PSP (2.21±0.12, *p*<0.001 compared to controls) whereas all other disease groups were not significantly higher compared to the controls (Fig. 1B). Calculation of the total vessel score (BG+FC) revealed PSP with the greatest total vessel score across neurodegenerative proteinopathies (PSP= 4.59±0.19, control = 2.36±0.32; *p*<0.0001) (Fig. 1C). MSA also showed significantly higher mean total vessel score (4.29±0.35, *p*<0.01) compared to controls. As many of these diseases present co-pathologies, we performed multiple logistic regression to assess the association between high vessel score and age, sex, and the presence of AD, LBD, MSA, PSP, AGD, and TDP-43 pathology, irrespective of the primary neuropathological diagnosis. Presence of PSP pathology among all conferred the highest odds ratio for overall high vessel pathology (OR = 6.820 for total vessel score ≥ 4, p < 0.001; Table 2), confirming the pronounced involvement of vessel wall thickening pathology in PSP across neurodegenerative proteinopathies.

**Figure 1.**
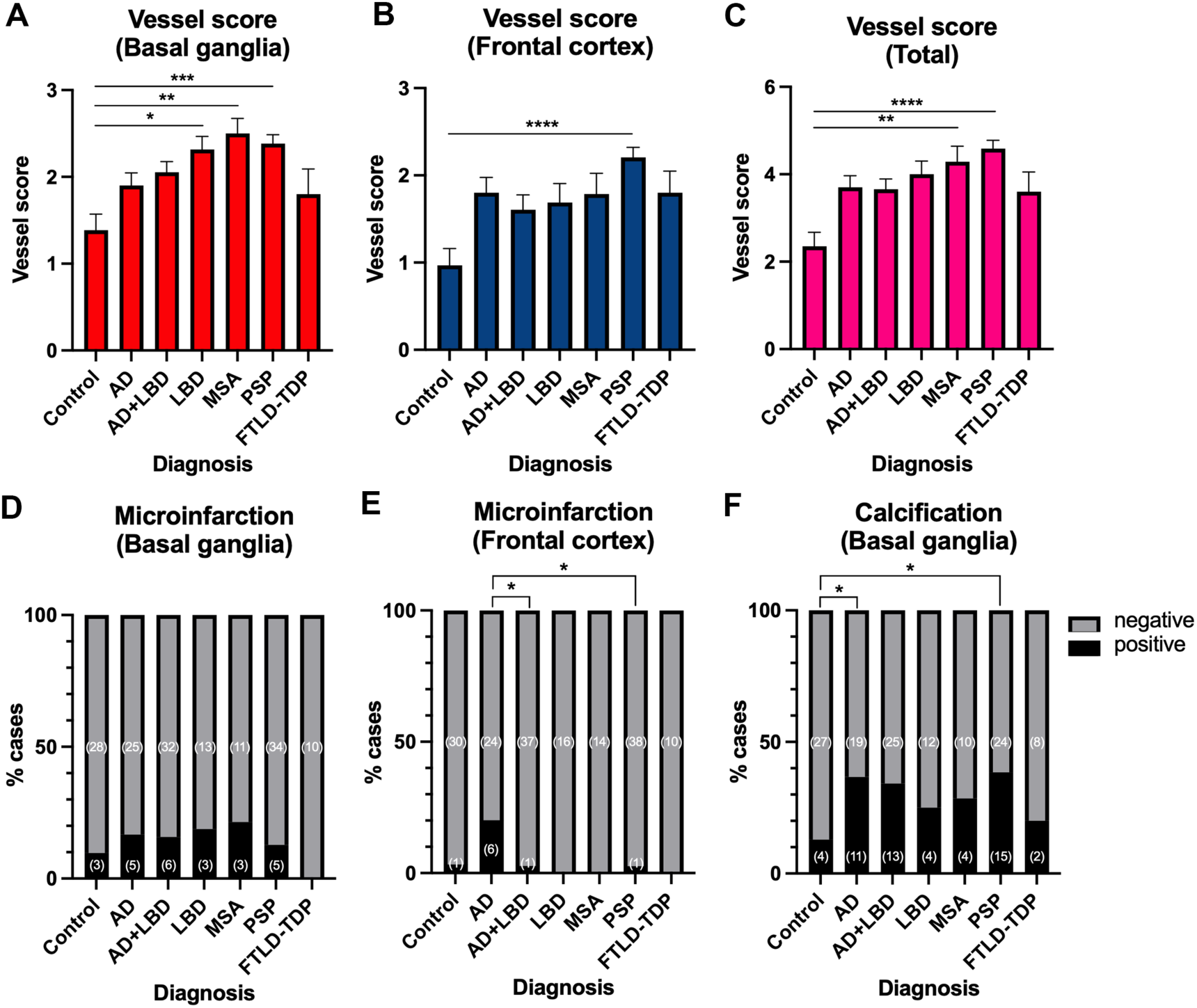
Vascular pathology in neurodegenerative disease groups. (A) Mean of vessel scores for the BG, (B) FC, and (C) total, along with the (D) frequency of microinfarcts in the BG and (E) FC, and (F) calcification in the BG were compared across control and disease groups. In the BG, MSA, PSP, and LBD in order showed significantly higher vessel scores compared to controls (A). In the FC, only PSP showed significantly higher vessel score compared to controls (B). Mean of total vessel scores (BG+FC) was highest in PSP (*p*<0.0001 compared to controls). MSA also showed significantly higher total vessel score compared to controls. No significant differences were found between any two disease groups (C). Frequency of microinfarction in the BG was not significantly different across the evaluated groups (D). In the FC, AD showed significantly higher frequency of microinfarction compared to AD+LBD and PSP groups (E). Calcification in the BG were significantly higher in AD and PSP groups compared to controls (F). Number of cases positive and negative for presence of microinfarction and calcification are indicated in brackets. \**p*<0.05, \*\**p*<0.01, \*\*\**p*<0.001, \*\*\*\**p*<0.0001 in (A-C) Dunn’s corrected multiple comparisons test post Kruskall-Wallis and (D-F) Fisher’s exact test. Abbreviations: Alzheimer’s disease (AD); Lewy body disease (LBD); Multiple system atrophy (MSA); Progressive supranuclear palsy (PSP); Frontotemporal lobar degeneration with TPD-43-immunoreactive pathology (FTLD-TDP); Basal ganglia (BG); Frontal cortex (FC).

**Figure 2.**
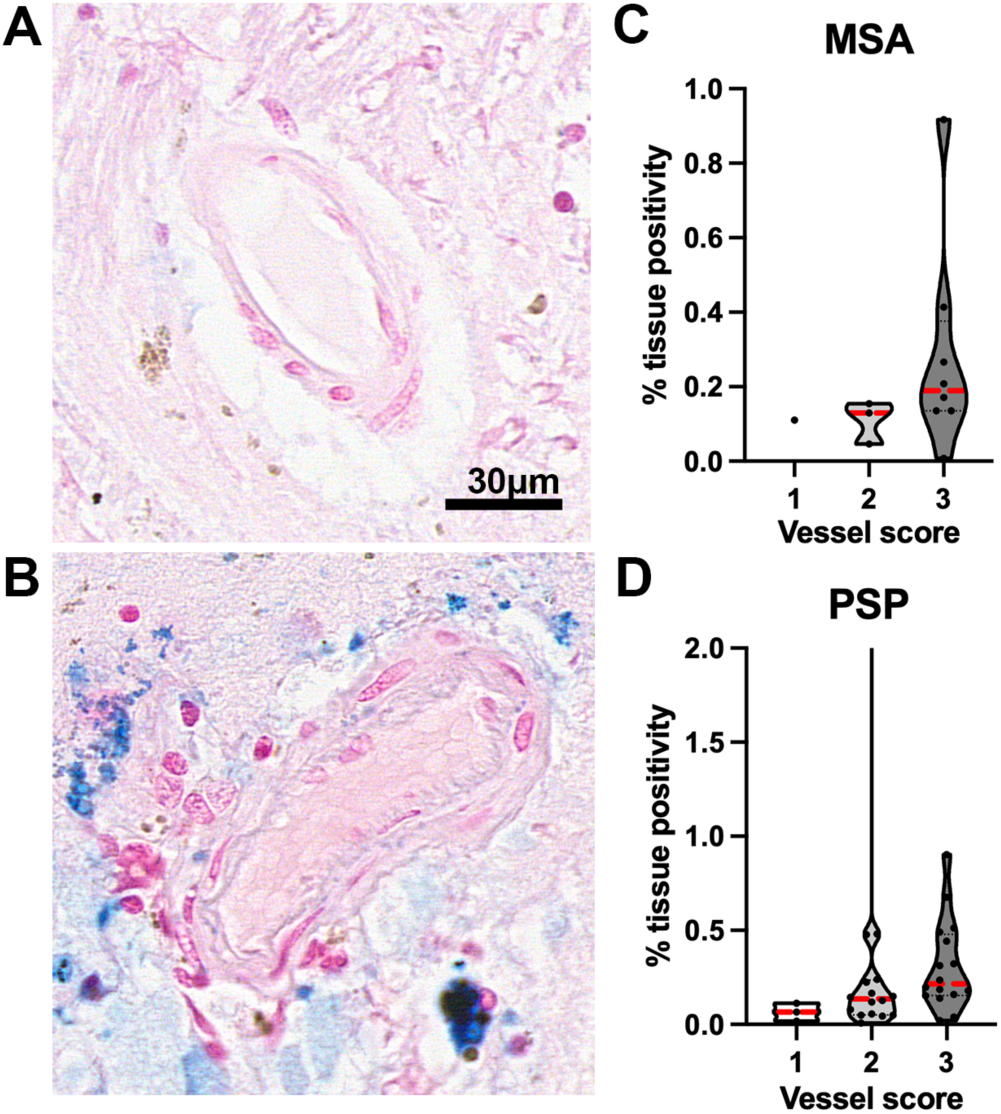
Iron accumulation associates with vessel scores in PSP and MSA. Blood vessels without (A) and with (B) perivascular parenchymal iron deposition visualized by Perl’s staining in the GP. (C) MSA cases with vessel score of 3 in the BG show greater GP iron levels as evaluated by % tissue positivity for Perl’s staining compared to those with vessel score 1 and 2. (D) In PSP, GP iron levels show an increasing trend with increasing BG vessel scores. One PSP case with BG vessel score of 2 shows extreme iron burden (% tissue positivity >5). Red dotted line represents the median and black dotted line represents the first and third quartile. Abbreviations: Multiple System Atrophy (MSA); Progressive Supranuclear Palsy (PSP); Globus pallidus (GP); Basal ganglia (BG).

**Table 2.**
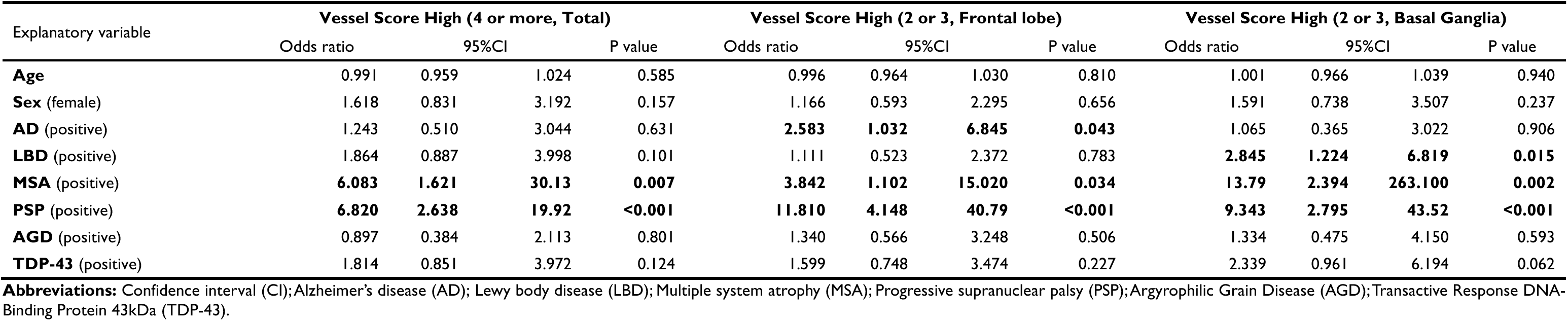
Regression Analysis for High Vessel Scores.

Compared to the observed disease-specific differences in the involvement of vessel wall thickening pathology, frequency of microinfarction in the BG did not significantly differ across all groups (Fig. 1D). In the FC, AD cases showed the greatest frequency of microinfarction (6/30 cases), that showed a significant difference compared to the lower frequencies in AD+LBD and PSP groups (1/37 and 1/38 cases, Fisher’s exact test; *p*<0.05 for both). (Fig. 1E). Equivalently lower frequencies are also found in other cohorts although the relatively lower number of cases in these groups may have limited the statistical significance. Calcification in the BG was significantly more frequent in AD (11/30 cases) and PSP (15/39 cases) groups compared to the controls (4/31 cases, *p*<0.05 for both) (Fig. 1F). However, multiple logistic regression analysis revealed no significant association between the presence of microinfarction nor calcification in the BG with any of the disease pathologies (Supplementary Table 1).

### Iron deposition associates with vessel scores in PSP and MSA

Given the significantly higher BG vessel scores observed in PSP and MSA—two disorders characterized by marked iron accumulation in the BG compared to other diseases—and considering that iron exchange occurs at the blood–brain barrier, we investigated the potential association between these two pathologies.

We quantified iron levels in the whole extent of GP (Fig. 2A and 2B) and evaluated the correlation with the BG vessel scores (Fig 2C and 2D). Analysis of Spearman’s *r* correlation revealed a significant correlation between GP iron load and vessel scores in PSP (*r*=0.424, *p*=0.011; n=35). MSA also showed a positive trend for correlation, which relationship may possibly be limited by case numbers (*r*=0. 521, *p*=0.085; n=12). Despite the reported correlation in PSP, one case with BG vessel score of 2 showed highest iron deposition amongst all cases (% tissue positivity >5) (Fig. 2D). These altogether suggests that although vessel wall thickening pathology may not be the sole contributor to iron deposition or vice versa in MSA and PSP, there is a general association between the two pathologies, particularly in PSP.

### Association of Vascular Pathology with *APOE* Genotypes and *MAPT* H1/H1 Haplotype

For possible contribution of genetic risk factors on the vessel pathologies of neurodegenerative disease brains, we evaluated the association of high vessel scores, BG microinfarcts and calcification with *APOE ε4* and *MAPT* H1/H1 haplotype positivity regardless of neuropathological diagnosis of the cases (Table 3). Neither *APOE ε4* nor *MAPT* H1/H1 haplotype were associated with vessel scores in the BG and the FC. *MAPT* H1/H1 haplotype positivity was inversely associated with microinfarction in the BG (OR=0.249, *p*=0.026). However, this association was attenuated after accounting for the protein pathologies in cases, suggesting that genetic risk factors are unlikely to substantially contribute to vascular pathology in neurodegenerative proteinopathies.

**Table 3.**
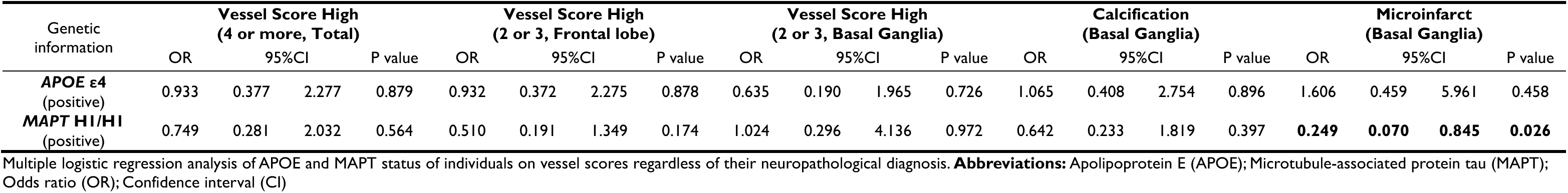
*APOE* and *MAPT* status on Microvascular Changes.

### SnRNA-seq reveals vascular cell proteotoxic stress associated with vessel wall thickening in PSP

Our histological findings highlighting the prominent association of vessel wall thickening in PSP across neurodegenerative proteinopathies prompted an investigation into the possible underlying mechanisms by snRNA-seq of vascular cell population (VC) isolated from PSP and control FC (Fig. 3A, Supplementary Table 2). Control cases with vessel scores in the FC of zero (n=4) and one (n=1), and PSP cases with scores zero (n=1), one (n=2), and two (n=5) were analyzed. After quality control and filtering of isolated nuclei and annotation of clusters, we obtained 0.71% of total nuclei to be of VC (Fig. 3B and 3C), which falls within previously reported vascular cell fraction obtained via sorting- and enrichment-free nuclei isolation methodologies of human post-mortem neurodegenerative brains, consistent with the method used here.^56–58^ Nonendothelial cell types that are not evaluated here are being parallelly examined in a separate study (Tanikawa *et al*.).^36^ Between the groups, the control group showed lower vascular fraction compared to PSP (0.55±0.15% and 1.54±0.34% respectively), however the endothelial cell/ pericyte composition within vascular fraction were similar between the groups (Fig. 3C). To validate cell type annotation of nuclei used for downstream analysis, the top 10 differentially expressed cell marker genes of each cell population were plotted against each other. (Fig. 3D) Identified endothelial markers included canonical and previously reported genes such as *EBF1*, *CLDN5*, *MECOM*, and VLMC-specific genes including *NOTCH3* were identified as pericyte markers.^59,60^

**Figure 3.**
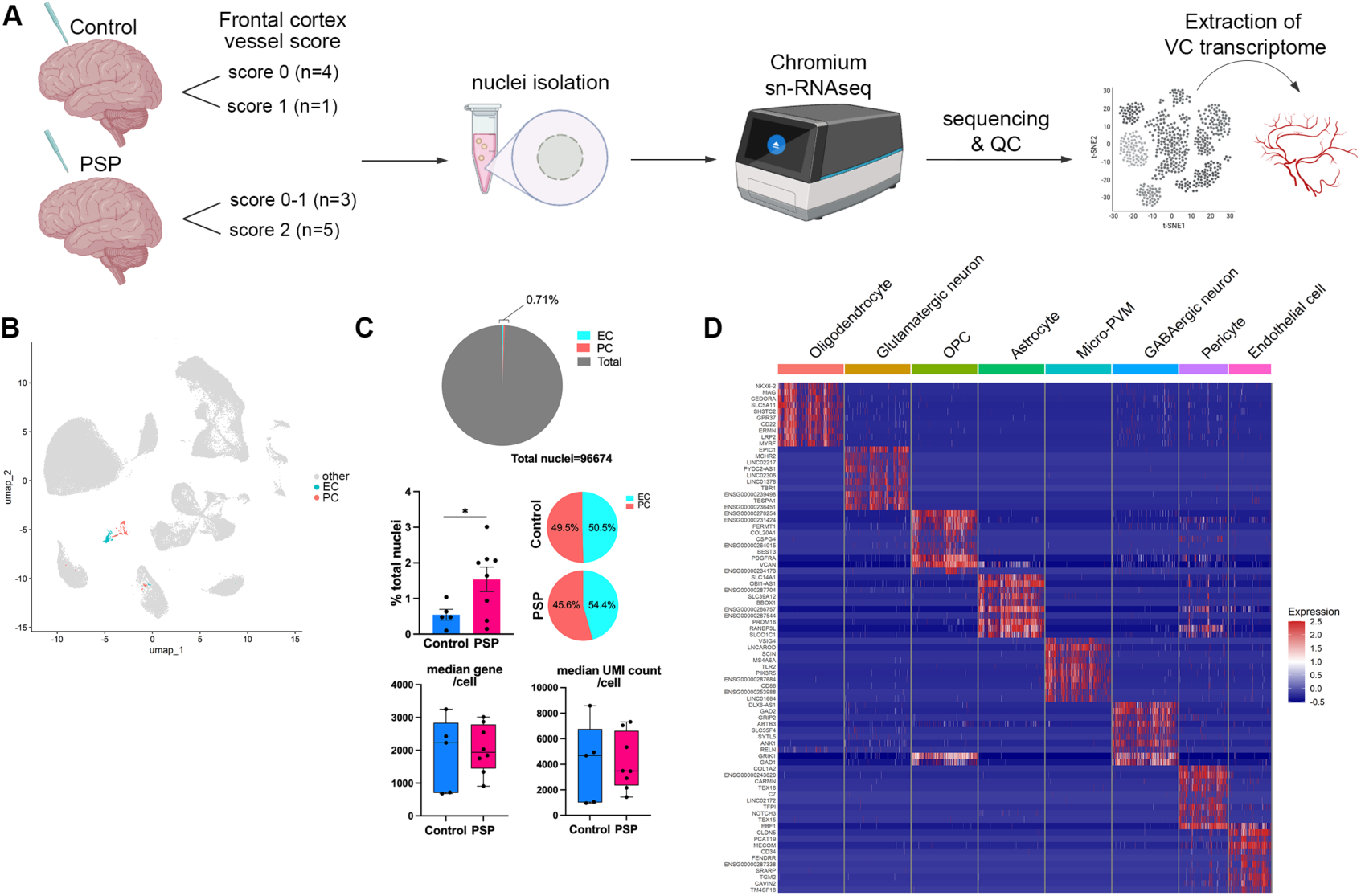
Single-nucleus RNA sequencing quality control and processing of control and PSP brain vasculature. (A) Schematic of snRNA-seq experiment. Frozen frontal cortex tissue of five control brains of vessel scores 0 and 1, eight PSP brains of vessel scores 0 to 2 were isolated for nuclei. Sequenced data were used to extract transcriptomes of endothelial cell (EC) and pericyte (PC) nuclei. (B) UMAP visualization of all isolated nuclei post quality control, among which EC are highlighted in cyan and PC in orange. (C) Quality control metrics for extracted EC and PC. 0.71% of total nuclei were of EC and PC. Vascular cell fraction was lower in the control group (mean ± SEM = 0.55 ± 0.15%) compared to PSP (1.54 ± 0.34%, p<0.05 by Student’s t-test). However, the relative composition of EC and PC remained similar between the groups. Median gene and UMI count per cell for all cases were plotted in box-and-whisker plots. (D) Top 10 cell type–specific marker genes for each cell population were plotted in heatmap for validation of cell type annotation, which include well-known EC and PC markers such as *CLDN5*, *EBF1*, and *CD34*. Abbreviations: Progressive supranuclear palsy (PSP); single nucleus RNA sequencing (snRNA-seq); quality control (QC); vascular cell (VC); oligodendrocyte precursor cell (OPC); endothelial cell (EC); pericyte (PC); perivascular macrophage (PVM).

Differential gene expression (DEG) analysis of endothelial transcriptome between control and PSP brains highlighted significant up-regulation of heat shock proteins in PSP which are believed to be up-regulated under stress: *HSP90AA1*, *HSPA1SA*, *HSPH1*, *HSPB1*, and *PTGES3* (Fig. 3E, Supplementary Table 3). Many of the significantly up-regulated genes were also of RNA-binding transcriptional and translational regulators (*HNRNPU*, *FXR1*, *TALAM1/MALAT1*) that are known to function in cellular stress-response.^61–63^ Interestingly, we found a significant up-regulation of *P4HA1* gene in PSP, a key enzyme in collagen synthesis which constitutes the extracellular matrix (ECM), and which has been shown to be involved in regulation of vascular wall remodeling^64,65^ – in alignment with our histological observations. On the other hand, mitochondrial genes vital for oxidative phosphorylation (*MT-ATP8*, *MT-ND3*) were significantly down-regulated. GOBP pathway enrichment analysis emphasized the up-regulation of protein folding and refolding-related pathways in PSP vasculature and revealed downregulation of cellular response mechanisms to ion and inorganic compounds (Fig. 4A). Interestingly, similar changes were seen in PSP pericytes, in which up-regulated genes were uniformly those involved in proteostasis stress response, including heat shock proteins (*HSPH1*, *HSPA1A*, *ST13*), *NR3C1,* coding for glucocorticoid receptor, and *SEM1*, coding for a structural subunit of 25S proteasome lid complex (Fig. 4A, Supplementary Table 4).

**Figure 4.**
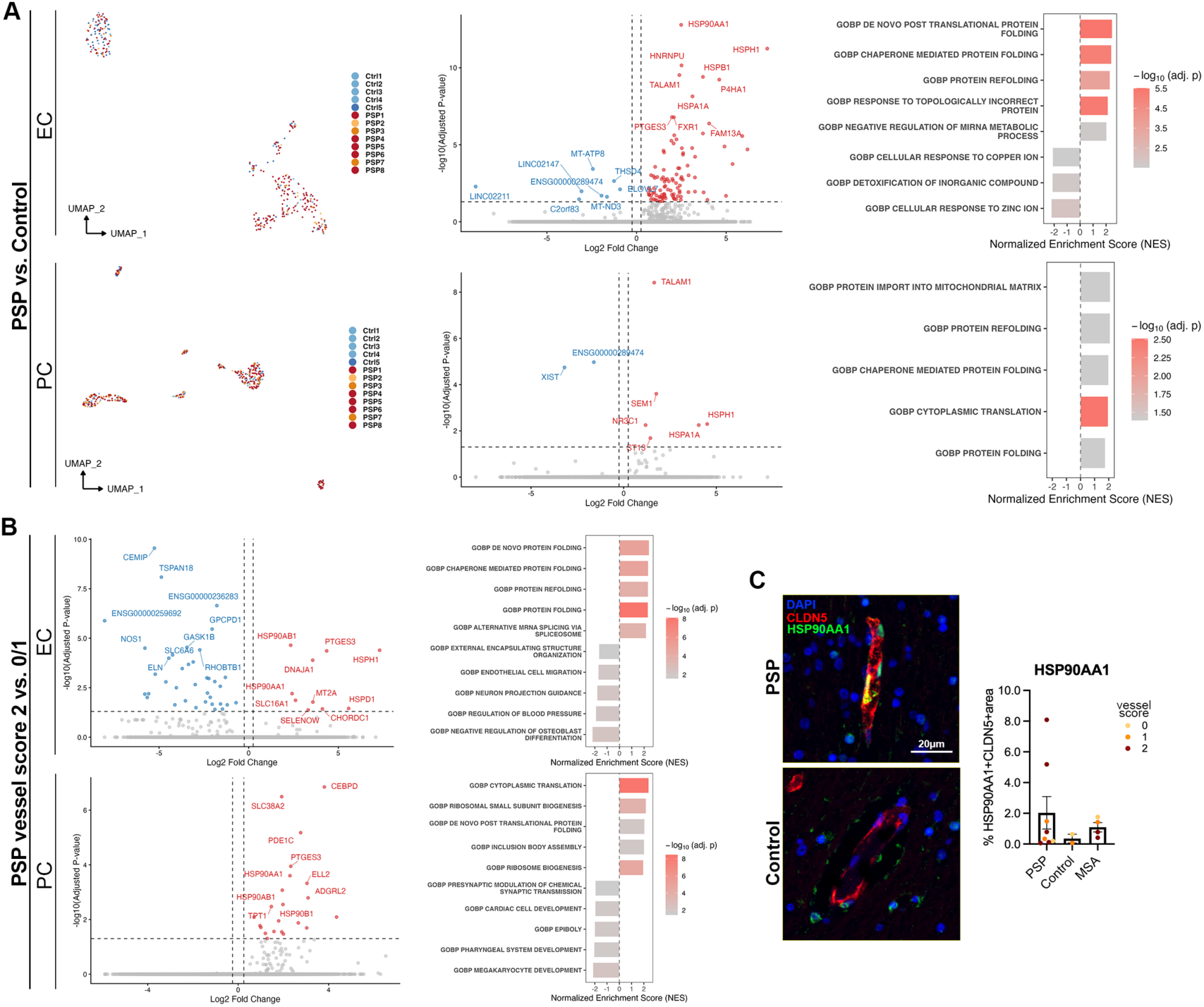
PSP EC and PC show proteotoxic signature. (A) UMAP visualization of EC and PC annotated by case. Control cases are annotated in increasing hue of blue by increasing vessel score (0 vs. 1), and PSP cases in increasing hue of red (0 vs. 1 vs. 2). Volcano plots showing significantly dysregulated EC and PC genes (adj. p-value <0.05) in PSP vs. control. GOBP pathway enrichment analyses highlight up-regulation of proteostasis-related pathways. (B) Evaluation of EC and PC transcriptome changes associated with higher vessel score within PSP cohort. Volcano plots showing significantly dysregulated EC and PC genes (adj. p-value <0.05) in PSP brains with vessel score 2 compared to those with scores 0 and 1 again highlight dysregulation of heat shock proteins. GOBP pathway enrichment analysis identifies relevant processes. (C) Double immunofluorescence of endothelial marker CLDN5 and heat shock protein HSP90AA1 for validation of its increased expression in PSP. Average endothelial HSP90AA1 expression quantified by %CLND5 area positive for HSP90AA1 (%HSP90AA1+CLDN5+area) is highest in PSP cases, followed by MSA then controls. Abbreviations: Progressive supranuclear palsy (PSP); endothelial cell (EC); pericyte (PC); claudin-5 (CLDN5); multiple system atrophy (MSA).

Next, given the heterogeneity in vessel scores within the PSP cohort, we examined the vascular transcriptome associated with higher vessel scores in PSP brains by stratifying cases into low (score 0 and 1) and high (score 2) vessel score group (Fig. 4B). Strikingly, DEG and pathway enrichment analysis of endothelial cells again primarily associated up-regulation of heat shock proteins and their chaperones, and protein folding and refolding pathways, with higher vessel scores (Fig. 4B, Supplementary Table 5). This suggests that the differences in PSP vascular transcriptome compared to controls are largely attributable to the observed vessel wall thickening pathology in the diseased brains. On the other hand, mainly genes involved in vasculature and ECM maintenance and remodelling were found to be downregulated in the higher vessel score group. Pericyte transcriptome revealed up-regulation of heat shock proteins and inflammatory markers (Fig. 4B, Supplementary Table 6).

We validated the expression of endothelial HSP90AA1 protein in the same PSP brains that were assessed for snRNA-seq using double immunolabelling with endothelial marker claudin-5 (Fig. 4C). We further evaluated their expression in cases with MSA, which we have previously shown to involve milder vessel wall thickening in the FC compared to PSP. In align with the vessel wall thickening scores across groups, we found the average expression levels of endothelial HSP90AA1, quantified by % CLDN5 area positive for HSP90AA1, were highest in PSP, followed by MSA then control, although the difference did not reach significance (Fig. 4C).

We present here a unique dataset of morphology-matched transcriptome of PSP and control human post-mortem brains but acknowledge the limitation of the small number of analyzed nuclei. Therefore, we sought to validate our findings in a larger independent snRNA-seq experiment analyzing in total 40 PSP, PiD, AD, and controls brains, from which the transcriptome of non-vascular nuclei was described by Rexach *et al*.^51^ (Fig. 5). Total of 8,085 endothelial cells from the insula, precentral gyrus, and the calcarine cortex were annotated and analyzed. First, we confirmed the integration of endothelial nuclei from the two datasets (Fig. 5A). Nuclei from the two datasets did not independently cluster, confirming comparable representation of endothelial cells in the analyzed nuclei of the two datasets. DEG analysis of each disease group compared to controls revealed distinct molecular signatures in PSP, PiD, and AD-associated endothelial cells (Fig. 5B) Across diseases, PSP endothelium was distinguished by the greatest dysregulation profile from controls, in alignment with its pronounced alteration observed at the histological level across diseases. Importantly, up-regulation of proteostasis-related genes in PSP endothelial cells were validated in this dataset, constituting the majority of significantly up-regulated DEGs (*HSPA1A*, *HSPA1B*, *HSPB1*, *CRYAB*, *HSPH1*, *PPID*), and further highlighted its proteotoxic signature in comparison to AD – where vessel wall thickening was insignificant compared to controls. PiD endothelial cells also demonstrated an up-regulation of heat shock proteins similar to that in PSP. In comparison to the previous dataset, cytokine-signalling associated genes were much heavily represented in the PSP down-regulated genes. Gene set enrichment analysis confirmed up-regulation of protein folding and refolding, and proteotoxic stress pathways in PSP that is not seen in AD (Fig. 5C). AD on the other hand showed up-regulation of pathways involved in endoplasmic reticulum and cell cycle. Again, PiD showed up-regulation of protein folding pathways alike PSP but PSP endothelial cell showed distinct down-regulation of cytokine signalling and vascular development and remodelling pathways. Since the dysregulated genes in this dataset were identified across three different brain regions, we next checked the genes involved in the top dysregulated – proteostasis and TNF signalling – pathways in each region. In PSP, genes of heat shock proteins and co-chaperones showed a consistent pattern of up-regulation compared to controls in each brain regions (Fig. 5D), which together with the previous FC dataset, support our argument from histological evaluation that vessel wall thickening pathology is a relatively global pathophysiology in PSP. Lastly, we identified TF regulons that are commonly PSP- and control-enriched in the two datasets for insight into key up-stream players of the disease-associated transcriptome alterations (Fig. 5E). Identified TFs included those involved in proteotoxic stress and suggested an overall poor stress-response state of PSP endothelial cells. Functional validation of these players awaits future experiments.

**Figure 5.**
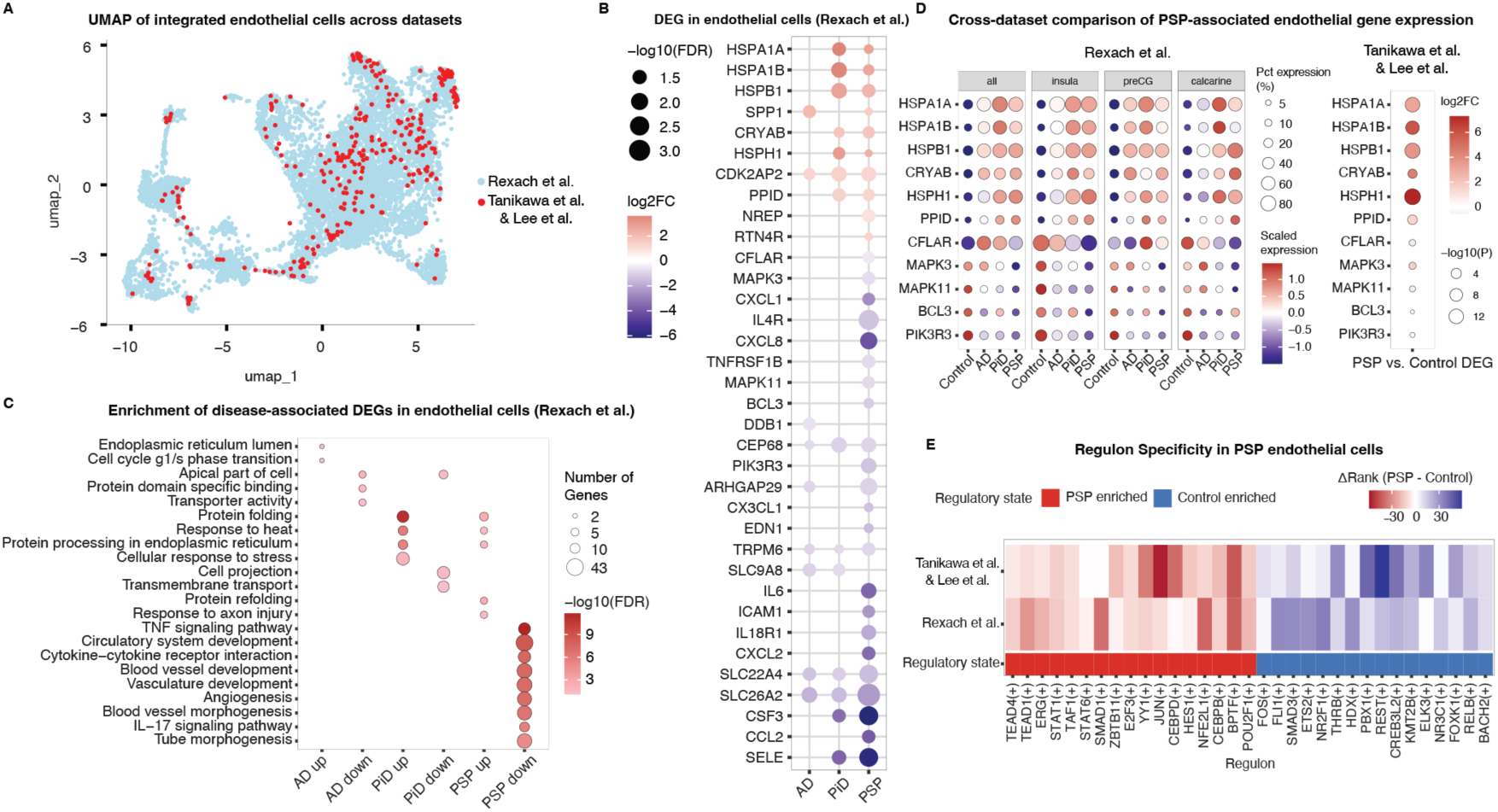
Shared PSP-associated transcriptional and regulatory signatures in endothelial cells across independent datasets. (A) UMAP visualization of endothelial cells integrated from two independent datasets (Rexach *et al*. and Tanikawa *et al*. & Lee *et al.*), colored by dataset, showing successful alignment across studies. (B) Differential expression analysis of endothelial cells in Rexach *et al.* across disease conditions (AD, PiD, PSP). Color indicates log2 fold-change relative to control, and dot size represents statistical significance. (C) Gene set enrichment analysis of differentially expressed genes in endothelial cells from Rexach *et al*., highlighting distinct pathways associated with diseases. Stress response and protein folding pathways are again prominent in PSP, with similar signature seen in PiD though PSP is distinctly characterized by significant down-regulation of inflammatory signaling and vascular remodeling pathways. AD show very different transcriptome from PiD and PSP. Dot size indicates the number of genes, and color represents significance (−log10 FDR). (D) Cross-dataset comparison of PSP-associated endothelial gene expression. Left: scaled single-cell normalized expression across brain regions and disease conditions in Rexach *et al.* Right: donor-level pseudobulk differential expression (log2 fold-change, PSP vs control) in Tanikawa *et al*. & Lee *et al.*, estimated using edgeR. (E) Cross-dataset comparison of regulon specificity for SCENIC-derived TF regulons. Regulon activity was quantified with AUCell, and specificity was assessed with the regulon specificity score (RSS). Within each dataset, regulons were ranked by RSS within each diagnosis (rank 1 indicates highest specificity). The difference in RSS rank between PSP and control (Δrank = rank_psp_− rank_control_) was used to quantify disease enrichment. Negative Δrank values (red) indicate higher specificity in PSP, whereas positive values (blue) indicate higher specificity in control. Regulons showing a consistent enrichment direction across both datasets were classified as shared, with PSP-enriched regulons shown in red and control-enriched regulons shown in blue.

## Discussion

Contribution of cerebrovascular pathologies to the pathogenesis of neurodegenerative proteinopathies is unclear. Our study evaluated the severity of brain arteriolosclerosis across multiple neurodegenerative proteinopathies and found that PSP and MSA exhibit severe changes in brain regions considered strategic for the development of protein pathology compared to other neurodegenerative diseases. We further identified local iron burden as a possible associating disease pathology in both disorders. SnRNA-seq analysis of the PSP vascular transcriptome across two independent cohorts revealed a prominent involvement of proteotoxic stress pathways in endothelial cells and pericytes. Importantly, this was associated with greater severity of vessel wall thickening pathology within the PSP cohort. Altogether our findings support microvasculature-associated pathomechanism as a possible contributor PSP.

Our neuropathological evaluation of SVD pathologies that are difficult to evaluate with patient imaging revealed differential involvement of microvascular alterations across neurodegenerative diseases. Notably, utilizing a scoring scheme adapted from VCING, we identified differing involvement of vessel wall thickening pathology across diseases, highlighting PSP for its significantly pronounced alteration in both BG and FC compared to controls, while MSA and LBD showed significant changes restricted to the BG (Fig. 1). The apparent vulnerability of the BG in PSP and MSA aligns with known patterns of early tau and α-syn pathology in these regions^30,66^, implicating vascular impairment in early disease pathogenesis. The more global involvement of vessel wall thickening in PSP may reflect differences in disease pathogenesis such as younger age of onset in MSA (50-60 years of age) compared to PSP (∼65 years).^67–69^ However, our regression analysis confirmed that high vessel scores were not associated with age, *APOE*, or *MAPT* risk factors, but were most strongly predicted by PSP followed by MSA pathology (Table 2 and 3) indicating that vessel wall thickening reflects a disease-specific pathophysiological process rather than a purely age-dependent or genetically mediated phenomenon.

Microvessel wall thickening reflects a structural and functional impairment of the cAVU.^70^ Key molecular mechanisms that underlie dysfunction in this unit include ECM remodeling, endothelial cell dysfunction and senescence, oxidative stress, and chronic inflammation,^71^ which are pathophysiologies strongly associated with early PSP and MSA. First, PSP tau pathology is observed to start accumulating in the subcortical regions including the BG, sites which are selectively vulnerable to aging-related subclinical hypoxia.^30,72^ In these regions, PSP brains show relevant changes, including ischemic lesions, white matter loosening, interruption of cellular respiration, increased intracellular iron deposition, and GP neuronal loss.^19,73^ Importantly, these areas and pathologies overlap with that seen in carbon monoxide toxicity.^74^ Next, early sites of PSP pathology uniquely accumulate excessive levels of iron that may underlie tau aggregation and neuroinflammation by the production of reactive oxygen species.^19^ Importantly, we found abnormal iron-laden astrocytes heavily localized adjacent to the vasculature, which together with a previous observation of perivascular orientation of early PSP astrocytic tau pathology,^21^ suggest that the disruption of the blood-brain-barrier (BBB) may be central to early PSP pathogenesis. More recently, the findings of pronounced CD8+ T-cell lymphocyte infiltration and microglial activation in the substantia nigra of PSP compared to PD, MSA, and aged controls evidence the involvement of heightened neuroinflammation and BBB permeability in PSP.^75,76^ Beyond this and together with the identification of a Human Leukocyte Antigen (HLA)-driven neuroinflammatory response^77^ in PSP, our findings highlight possible avenues for stratification for preventive strategies. In comparison, early-effected striatonigral regions of MSA brains show activated microglia, iron accumulation, and up-regulation of chronic hypoxic marker, HIF-2α.^35,78^ Notably, case reports of cerebrovascular lesions without significant protein pathology at autopsy, presenting with MSA-like autonomic and motor symptoms, suggest that early autonomic dysfunction in MSA may reflect vascular impairment preceding or independent of canonical proteinopathy.^78–80^ We further explored the association between microvascular alterations and iron deposition in the BG of PSP and MSA, identifying a possible pathomechanistic link that may contribute to tau and α-syn accumulation as described in more detail in this review^81^ (Fig. 2). However, the directionality of identified relationships remains uncertain: vascular impairment may drive protein aggregation, or conversely, tau and α-syn deposition may itself trigger vascular damage, establishing a vicious cycle of BBB dysfunction. The precise temporal relationship between protein pathology and vascular impairment in PSP and MSA therefore warrants further investigation.

Our snRNA-seq analysis of PSP and control vascular population revealed a strong and uniform involvement of proteotoxic stress in PSP vasculature (Fig. 3 and 4), in contrast to similar studies highlighting angiogenesis-related changes in AD.^58,59,82–84^ This finding was validated across analyses of two independent PSP cohorts and four anatomical regions. The preserved signature across brain regions supports our histological conclusions that vessel wall thickening is a global pathophysiology of PSP. Importantly, our datasets included endothelial nuclei from AD and PiD brains. While we did not detect a strong angiogenic change reported previously in AD, we confirmed markedly distinct endothelial transcriptomes; notably, a stressed proteostasis signature was absent in AD endothelial cells (Fig. 5). These transcriptomic differences align with our histological findings, where microinfarctions were more frequent in AD than vessel wall thickening (Fig. 1), suggesting distinct vascular impairments or pathomechanisms between the two diseases. Interestingly, we found proteotoxic stress to be a shared molecular signature in PiD endothelia. Although histological scoring of vessel wall thickening in PiD brains were not pursued in our original cohort, as mentioned previously, a study showed more widespread SVD in PiD brains compared to AD, PSP, AGD, and CBD.^23^ Future study comparing regional and global severity of specific microvascular pathologies and their molecular signatures across FTLD-tau may reveal possible overlapping or distinct pathomechanisms.

In parallel to proteostasis stress-related genes, up-regulated genes in PSP endothelial cells included many RNA-binding proteins (*HNRNPU*, *TALAM1*, *FXR1*, *LINC0221*, *LINC01068*, *MCF2L-AS1*, and *NREP*) which are frequently reported to be dysregulated with aging.^85,86^ Some of these (*HNRNPU*, *TALAM1*/*MALAT1*, *FXR1)*, and others (*P4HA1*) are reported to respond to cellular stress including hypoxia.^61–63,65^ Accumulation of misfolded proteins and induction of heat shock protein expression have been directly connected to hypoxia,^87,88^ which further supports the notion that hypoxia may be an underlying or preceding mechanism to the pathological endothelial transcriptome observed in PSP. Among hypoxia-response genes, *P4HA1*, which is transcriptionally activated by HIF-α, has been implicated in fibrosis and atherosclerotic plaque formation^65^, suggesting a potential role in disease-associated vessel wall thickening. However, within the PSP cohort, upregulation of heat shock proteins showed a stronger association with higher vessel scores than *P4HA1*, pointing to proteostasis stress as a more prominent molecular correlate of vascular pathology in PSP.

Our PSP findings are further congruent with a prior transcriptomics study examining superior temporal cortex, which reported a significant up-regulation of stress response and immune system process in PSP microglial/endothelial population.^89^ They also found key genes *HSP90AA1* and *HSPB1* significantly up-regulated in pericytes, with additional markers *HSPH1*, *HSPA1A*, *PTGES3*, and *ST13* detected using unadjusted p-values (https://rtools.mayo.edu/PSP_RNAseq_Atlas/). The same study reported changes in the microglial/endothelial population to be one of most prominent disease-associated alterations, which supports vasculature dysfunction as a core feature of PSP. Notably, their findings in the relatively tau-sparing temporal cortex compared to the evaluated BG and FC here further support a global vascular involvement in PSP.

As a limitation of the present study, we were unable to correlate our findings with established cardiovascular risk factors, including diabetes mellitus, arterial hypertension, and hypercholesterolemia, which are known to contribute to disease-associated microvascular alterations. However, we did not detect fibrinoid necrosis, and these risk factors are associated with more severe lesions.^90^ Furthermore, PSP and MSA lack evidenced association with cardiovascular risk factors, rendering the association between the pronounced arteriolosclerosis we identify here with such risk factors less likely.^91^ To note, a multicentre epidemiology study had identified hypertension to be significantly associated with PSP,^92^ suggesting a possible contribution to the observed global vessel wall alterations. However, again hypertension is more closely associated with fibrinoid necrosis of the vessel walls. Importantly, we show that the endothelial signature in AD, which is also known to associate with hypertension^93–95^, is different from that in PSP suggesting the observed vessel wall thickening in PSP is underlined by different mechanisms.

*In vitro*, endothelial cells have shown to internalize exogenous tau fibrils^96^ and in AD and PSP brains, pathogenic tau has also been reported to accumulate within the cerebral microvasculature. One study claimed T22 antibody positivity in vessel walls^97^; these findings would benefit from confirmation using additional antibodies. Another reported diffuse, vessel-associated pattern of phosphorylated tau accumulation along the surface of select blood vessels in AD.^98^ Further, one study has reported prominent up-regulation of heat shock protein transcripts in snRNA-seq analysis of human AD brain^60^ which altogether may alternatively suggest a general tau-induced mechanism of endothelial stress signature. Regardless, our histological evaluation reveals clear differences in the regional severity and type of vessel pathology across disease (Fig. 1) supporting distinct pathomechanisms in disease. Importantly, we show consistent up-regulation of stress response protein transcripts and down-regulation of genes involved in ECM maintenance in PSP cases with high vessel score compared to low (Fig. 4B), providing a morphological correlate to the neurodegenerative endothelial and pericyte transcriptome associated with PSP tau pathology. Future PSP studies utilizing isolation methodologies for enrichment of vasculature nuclei^99^ that will allow comprehensive evaluation by origin of zonation will strengthen our findings on microvasculature morphology-transcriptome correlation.

In conclusion, our neuropathological analysis across neurodegenerative proteinopathies identifies vessel wall thickening with elevated endothelial and pericyte proteotoxic stress as a prominent feature of PSP. While the spatiotemporal relationship with tau pathology and its contribution to disease progression and clinical symptoms require further study, the disease specificity, potential early involvement, and widespread vascular alterations support microvasculature and cAVU dysfunction as plausible early therapeutic targets in PSP.

## Data availability

Raw sequencing data and annotated data objects will be made publicly available upon publication. Source data for all main and extended data figures will be provided with the paper. Custom code used for preprocessing and downstream analyses is available from the corresponding author upon reasonable request.

## Acknowledgments

The authors thank the patients who made this research possible. Brain samples were obtained through appropriate consenting procedures, with approval from the Local Ethic Committee and the University Health Network Research Ethics Board (Nr. 20–5258).

## Funding

This study was supported by the Edmond J Safra Philanthropic Foundation, the Rossy Family Foundation, Canadian Foundation for Innovation (40480), and the Ontario Research Fund. Gabor G. Kovacs holds the Rossy Chair in PSP Research.

## Competing interests

AEL has served as an advisor for AbbVie, Amylyx, Biogen, Biohaven, BlueRock, EG427, Ferrer, Janssen, and Ventyx.; received honoraria from AbbVie; received grants from Canadian Institutes of Health Research, Edmond J Safra Philanthropic Foundation, Krembil Brain Institute, Michael J. Fox Foundation, Parkinson Foundation, Parkinson Canada and The Rossy Foundation; has served as an expert witness in litigation related to paraquat and Parkinson’s disease, and has received publishing royalties from Elsevier, Saunders, Wiley-Blackwell, Johns Hopkins Press, and Cambridge University Press. ; GGK and AEL shared pending patent for Diagnostic assays for movement disorders (18/537,455). GGK has a shared patent for 5G4 synuclein antibody. GGK reports personal fees from Parexel, Mitsubishi-Tanabe, Ferrer, and Treventis; other funding from MSA Coalition, Michael J. Fox Foundation, Parkinson Canada, NIH, Canada Foundation for Innovation, and Ontario Research Fund; in addition, GGK received royalties from Wiley, Cambridge, Taylor & Francis, and Elsevier publishers. SLF is supported by the National Health and Medical Research Council of Australia Ideas grant (#214090508). MCT receives consultancy support from Biogen and Hoffman-La Roche and is a board member of Alzheimer’s Society of Toronto. SHF receives research Funding from Michael J. Fox Foundation for Parkinson Research, NIH (Dystonia Coalition); Parkinson Canada; Weston Foundation. Honoraria from the International Parkinson and Movement Disorder Society. Consultancy/Speaker fees from Abbvie, Lundbeck, Sunovion and Royalties from Oxford University Press. These have no relevance to this study. SL, XH, ST, TK, KY, SI, HT, TK, ST, ER and JER declare no competing interests.

## Consent statement

All brain specimens were approved by the UHN Research Ethics Committee (No. 20-5258) and the University of Toronto (No. 39459) and performed according to the ethical standards outlined in the 1964 Declaration of Helsinki (updated 2008).

## Supplementary material

Supplementary tables are provided in separate files.

