## Supplementary for "Microvascular pathology of proteotoxic endothelial signature characterizes Progressive Supranuclear Palsy"

### Methodology

#### Case selection

For disease characterization and staging, formalin-fixed paraffin-embedded (FFPE) tissue sections of 4.5 µm thickness were stained with hematoxylin, eosin, and luxol fast blue (HE/LFB), phosphorylated tau (pTau, clone AT8, 1:1000, Invitrogen/ThermoFisher, Carlsbad, USA), Aβ (clone 6F/3D, 1:50, Dako, Glostrup, Denmark), alpha-synuclein (a-syn) (clone 5G4, 1:4000, Analytikjena, Jena, Germany), and phosphorylated TDP-43 (pTDP-43, clone 11-9, 1:2000, CosmoBio, Tokyo, Japan) antibodies. Additionally, for pTau-positive cases, 3-repeat-tau (3R-tau, clone 8B6/C11, RD3, 1:5000, MilliporeSigma, Bedford, USA) and 4-repeat-tau (4R-tau, clone 1E1/A6, RD4, 1:200, MilliporeSigma) were stained. Following the manufacturer's instructions, immunostaining was carried out using Dako Autostainer Link 48 and EnVision FLEX+ Visualization System.

#### Single-nucleus RNA sequencing analysis

##### *Cell-type marker analysis and heatmap*

Cell type–specific marker genes were identified in Seurat. Positive markers for each cell type were computed using the FindAllMarkers function, and the top ten genes per cell type were selected based on average log2 fold change. To visualize these markers, up to 500 cells per cell type were randomly subsampled, scaled expression values were computed with ScaleData, and a heatmap of the selected genes was generated using DoHeatmap.

##### *Visualization of differential gene expression analysis*

Volcano plots were generated in RStudio using ggplot2 (v4.0.0)[1] and ggrepel (v0.9.6).[2] For each gene, the average $\log_2$ fold change returned by FindMarkers (avg_$\log_2$FC; PSP versus controls) was plotted on the x-axis, and $-\log_{10}$(p_val_adj) on the y-axis. Genes with p_val_adj < 0.05 and avg_$\log_2$FC > 0.25 were labeled as upregulated, those with p_val_adj < 0.05 and avg_$\log_2$FC < −0.25 as downregulated, and all others as not significant, and points were colored accordingly. The top 10 upregulated and top 10 downregulated genes (ranked by p_val_adj) were annotated on the plots.

***Summary of snRNA-seq data processing from Rexach et al.*** [3]

Briefly, raw snRNA-seq data were processed using Cell Ranger (v3.0) and aligned to the human reference genome (Ensembl release 93). Cells with <200 detected genes or >8% mitochondrial reads were excluded, and genes detected in fewer than three cells were removed. Expression values were normalized by library size (UMI counts), log-transformed, and corrected for sequencing depth and technical covariates; doublets and outlier cells were identified and removed prior to downstream analysis. Cell types were identified by joint clustering and annotated using canonical marker genes, with classifications further validated using reference-based mapping to external human brain datasets.

1. Wickham H. ggplot2. 2016.
2. Kamil Slowikowski TvdB, Alicia Schep, Sean Hughes, Trung Kien Dang, Saulius Lukauskas, Jean-Olivier Irisson, Zhian N Kamvar, Thompson Ryan, Dervieux Christophe, Yutani Hiroaki, Pierre Gramme, Amir Masoud Abdol, Malcolm Barrett, Robrecht Cannoodt, Michał Krassowski, Michael Chirico, Pedro Aphalo, Francis Barton. ggrepel: Automatically Position Non-Overlapping Text Labels with 'ggplot2'. doi:10.32614/CRAN.package.ggrepel
3. Rexach JE, Cheng Y, Chen L, et al. Cross-disorder and disease-specific pathways in dementia revealed by single-cell genomics. *Cell*. Oct 3 2024;187(20):5753–5774.e28. doi:10.1016/j.cell.2024.08.019

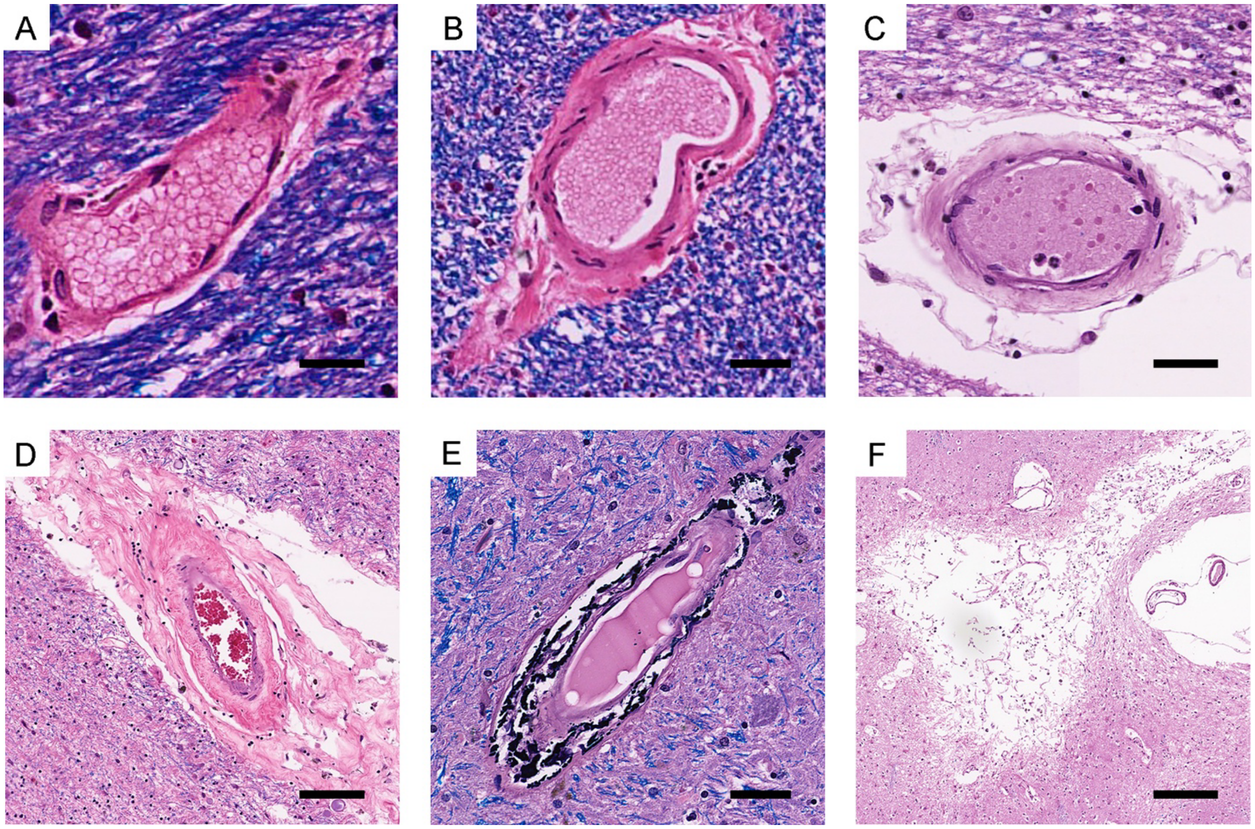


**Supplementary Figure 1.** Microscopic images of vascular pathology. Vessel wall thickening was graded according to the Vascular Cognitive Impairment Neuropathology Guidelines (VCING) as follows: (A) normal vessels were graded as grade 0, (B) vessels with wall thickening but no fatty hyaline deposits were graded as grade 1, (C) vessels with fatty hyaline deposits but no obvious luminal narrowing were graded as grade 2, and (D) vessels with obvious luminal narrowing were graded as grade 3. Additionally, (E) the presence of calcification and (F) microinfarction were assessed in both the basal ganglia and frontal lobe. Bars represent 15 μm for A and B, 25 μm for C, 50 μm for D and E, and 100 μm for F.

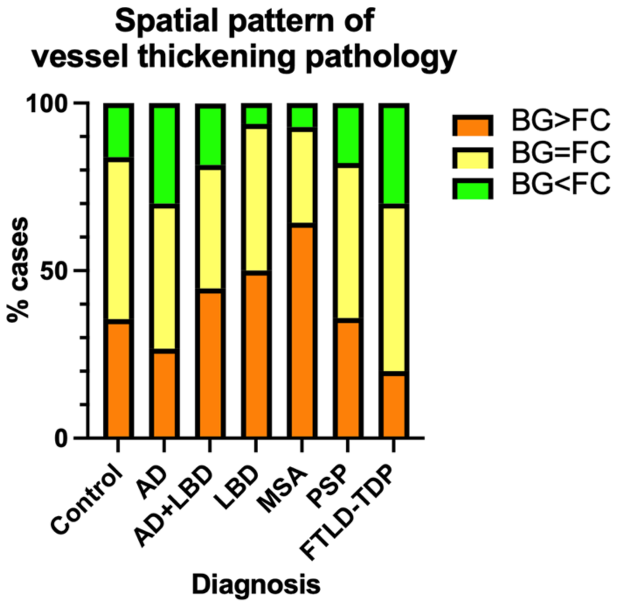


**Supplementary Figure 2.** Percentage of cases per disease group with a higher vessel score in the BG compared to the FC (orange), vice versa (green), and equal scores in both regions (yellow). Generally, vessel pathology followed the spatiotemporal pattern of protein pathology. MSA and LBD cohort showed greater percentage of cases with vessel scores higher in the BG compared to the FC, which pattern was followed to a lesser extent in the AD+LBD. In AD, comparably higher percentage of cases showed greater vessel score in the FC compared to the BG. In PSP, greater percentage of cases showed higher vessel scores in the BG than the FC, with generally similar patterns to that of the control cohort.

**Supplementary Table 1. Regression analysis for microinfarction and calcification in the basal ganglia**

| Explanatory variable (target side) | Microinfarction | | | | Calcification | | | |
|---|---|---|---|---|---|---|---|---|
| | Odds ratio | 95%CI | | P value | Odds ratio | 95%CI | | P value |
| **Age** | 1.032 | 0.984 | 1.087 | 0.199 | 1.025 | 0.990 | 1.064 | 0.168 |
| **Sex** (female) | 2.243 | 0.899 | 5.98 | 0.084 | 1.236 | 0.630 | 2.430 | 0.537 |
| **AD** (positive) | 2.082 | 0.553 | 10.51 | 0.293 | 1.311 | 0.544 | 3.297 | 0.551 |
| **LBD** (positive) | 1.270 | 0.464 | 3.593 | 0.643 | 0.862 | 0.405 | 1.832 | 0.699 |
| **MSA** (positive) | 3.701 | 0.652 | 18.39 | 0.131 | 1.309 | 0.316 | 4.726 | 0.693 |
| **PSP** (positive) | 1.326 | 0.377 | 4.204 | 0.644 | 1.735 | 0.743 | 4.041 | 0.201 |
| **AGD** (positive) | 0.481 | 0.126 | 1.841 | 0.212 | 0.737 | 0.312 | 1.662 | 0.528 |
| **TDP-43** (positive) | 1.385 | 0.497 | 3.700 | 0.524 | 1.070 | 0.483 | 2.312 | 0.866 |

**Abbreviations:** confidence interval (CI); Alzheimer's disease (AD); Lewy body disease (LBD); Multiple system atrophy (MSA); Progressive supranuclear palsy (PSP); Argyrophilic Grain Disease (AGD); TAR DNA-binding protein 43 (TDP-43).

**Supplementary Table 2. Summary of cases used in single-nuclear RNA sequencing**

| | Age, mean±SD (yrs) | Sex (F/M) | PMD, mean±SD (hrs) | ADNC, intermediate – high, *n* (%) | Lewy related pathology positive, *n* (%) | LATE-NC positive, *n* (%) | AGD positive, *n* (%) |
|---|---|---|---|---|---|---|---|
| **Control** | | | | | | | |
| FC vessel score 0 (*n*=4) | 69.3±11.6 | 1/3 | 12.7±10.8 | 0 (0.0) | 0 (0.0) | 0 (0.0) | 0 (0.0) |
| FC vessel score 1 (*n*=1) | 64.0±0.0 | 1/0 | 24.0±0.0 | 0 (0.0) | 0 (0.0) | 0 (0.0) | 0 (0.0) |
| **PSP** | | | | | | | |
| FC vessel score 0/1 (*n*=3) | 76.0±3.0 | 0/3 | 7.5±5.7 | 0 (0.0) | 0 (0.0) | 0 (0.0) | 1 (33.3) |
| FC vessel score 2 (*n*=5) | 73.4±3.0 | 1/4 | 21.5±19.5 | 0 (0.0) | 1 (20.0) | 0 (0.0) | 1 (20.0) |

**Abbreviations:** Frontal cortex (FC); Progressive supranuclear palsy (PSP); standard deviation (SD); post-mortem delay (PMD); Alzheimer's disease neuropathological change (ADNC); Limbic-predominant age-related TDP-43 encephalopathy neuropathologic change (LATE-NC); Argyrophilic Grain Disease (AGD).
